# Whole genome profiling of short-term hypoxia induced genes and identification of HIF-1 binding sites provide insights into HIF-1 function in *Caenorhabditis elegans*

**DOI:** 10.1101/2023.11.15.567310

**Authors:** Dingxia Feng, Long Qu

## Abstract

Oxygen is essential to all the aerobic organisms. However, during normal development, disease and homeostasis, organisms are often challenged by hypoxia (oxygen deprivation). Hypoxia-inducible transcription factors (HIFs) are master regulators of hypoxia response and are evolutionarily conserved in metazoans. The homolog of HIF in the genetic model organism *C. elegans* is HIF-1. In this study, we aimed to understand short-term hypoxia response and to identify HIF-1 direct targets in *C. elegans*. The central research questions were: (1) which genes are differentially expressed in response to short-term hypoxia? (2) Which of these changes in gene expression are dependent upon HIF-1 function? (3) How do HIF-1-dependent hypoxia-responsive genes affect hypoxia adaptation? (4) Which genes are the direct targets of HIF-1? We combine whole genome gene expression analyses and chromatin immunoprecipitation sequencing (ChIP-seq) experiments to address these questions. In agreement with other published studies, we report that HIF-1-dependent hypoxia-responsive genes are involved in metabolism, oxidation-reduction process, and stress response. Some HIF-1-dependent hypoxia-responsive genes like *efk-1* and *phy-2* dramatically impact survival in hypoxic conditions. HIF-1 co-immunoprecipitates with genomic regions proximal genes involved in stress response, protein processing in endoplasmic reticulum, and cell recognition. Further, some of these potential HIF-1 direct targets are differentially expressed under short-term hypoxia or are differentially regulated by mutations that enhance HIF-1 activity.

## Introduction

Oxygen is essential to aerobic organisms for energy production and cellular redox environment maintenance [1]. However, during development, disease and homeostasis, animals are often challenged by oxygen deprivation (hypoxia). In mammals, the majority of transcriptional response to hypoxia is mediated by the hypoxia-inducible transcription factors HIFs. Understanding HIF function and regulation has the potential therapeutic significance in common hypoxia-related diseases, such as cancer, arthritis and ischemia. The heterodimeric HIF complexes consist of α and β subunits, and both subunits are bHLH (basic-helix-loop-helix)-PAS (PER/ARNT/SIM) domain proteins [1, 2]. There are three HIFα and three HIFβ homologs in the human genome [3]. HIFβ has multiple bHLH-PAS dimerization partners and is relatively stable and abundant. In contrast, HIFα is short-lived under well-oxygenated conditions and is dedicated to hypoxia response [4–6]. The stability of mammalian HIFα is regulated by the PHD-VHL pathway. When oxygen levels are high enough, HIFα is hydroxylated by prolyl hydroxylase proteins (PHDs), in reactions that require oxygen as substrate. The hydroxylated HIFα is then targeted by the E3 ligase VHL (von Hippel-Lindau tumor suppressor) for proteasomal degradation.

The nematode *C. elegans* is a powerful genetic model organism for studying HIF function and regulation [7], and its genome encodes single homologs for HIFα and HIFβ, named HIF-1 and AHA-1, respectively. While *hif-1*α -/- mice die by E9.0 with severe vascular defects [8, 9], *C. elegans hif-1*(*ia04*) loss-of-function mutants survive and develop normally in room air but are defective in hypoxia adaption [10–12]. As in mammals, the AHA-1 subunit is stable, while the stability of HIF-1 is regulated by oxygen levels [11, 13, 14]. The PHD-VHL pathway for HIF-1 stability regulation is simplified in *C. elegans*, too. There is only one counterpart for PHD and VHL in *C. elegans*, named EGL-9 and VHL-1, respectively [13]. In addition to its role in targeting HIF-1 for oxygen-dependent degradation, EGL-9 also inhibits HIF-1 activity: HIF-1 targets are expressed at higher levels in *egl-9(sa307)* loss-of-function mutants than in *vhl-1(ok161)* loss-of-function mutants [15, 16].

Prior studies [17, 18] have described oxygen-dependent changes in *C. elegans* gene expression, but the direct targets of HIF-1 are not fully described. Here, we take advantage of *C. elegans egl-9* mutants that fail to degrade HIF-1 and ask which DNA sequences co-immunoprecipitate with HIF-1. To determine which of these targets are likely the immediate targets of HIF-1, we identify the genes that are differentially regulated by short-term (2 hours) hypoxia treatment and ask whether some of the HIF-1 targets are essential to survival in hypoxic conditions.

## Results

### Identification of genes responsive to short-term hypoxia treatment

To define the immediate gene expression changes caused by hypoxia, we treated L4-stage wild-type N2 worms with 0.5% oxygen for 2 hours and compared the whole genome gene expression profile under hypoxia with that in room air. The complete analysis results are provided in S1 Table. Among the 18,011 unique genes assayed, the mRNA levels of 681 genes were changed by hypoxia: 437 genes were up-regulated (S2 Table), and 244 genes were down-regulated (S3 Table). The enriched biological functions for genes up-regulated by short-term hypoxia were oxidation-reduction process, response to stress, transition metal ion binding, small molecule metabolic process, and cellular response to unfolded protein (Table 1). The representative genes for each functional group were listed in Table 1. The full lists of genes for each functional group were provided in S4 Table.

**Table 1.**
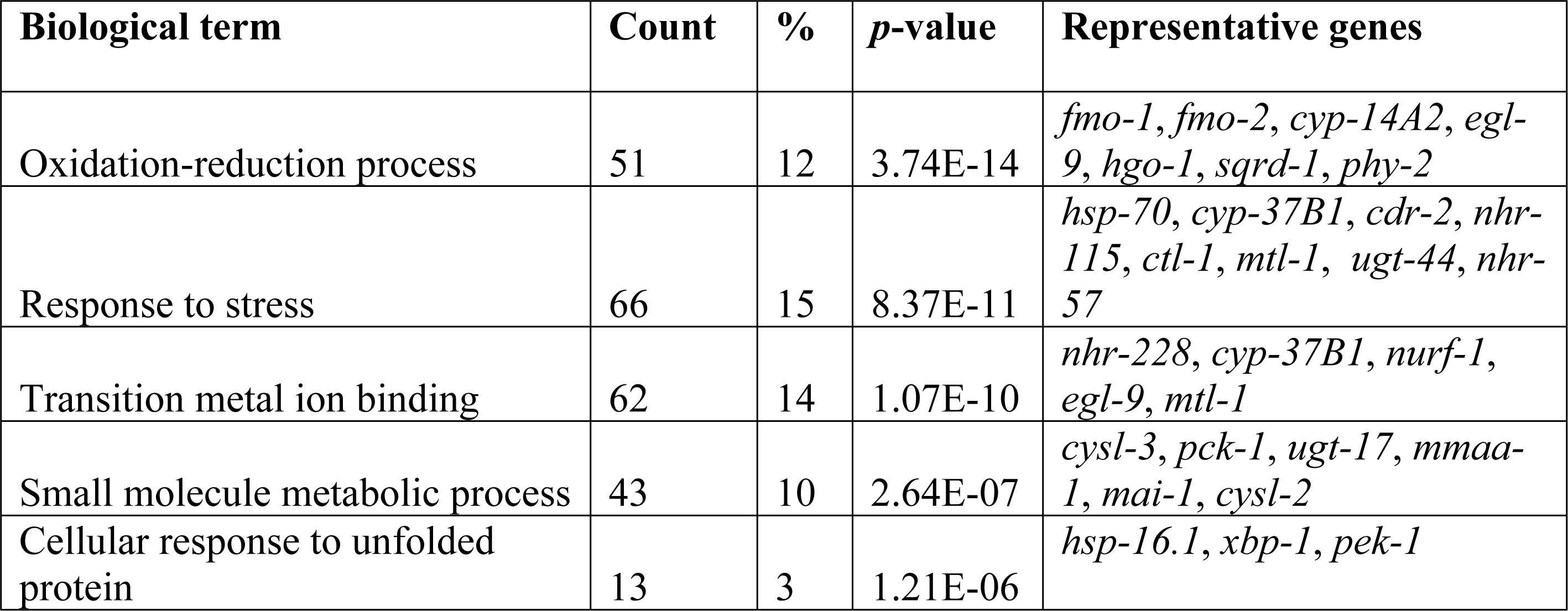
Enriched biological terms for genes up-regulated by short-term hypoxia in N2.

| Biological term | Count | % | *p*-value | Representative genes |
| --- | --- | --- | --- | --- |
| Oxidation-reduction process | 51 | 12 | 3.74E-14 | *fmo-1, fmo-2, cyp-14A2, egl-9, hgo-1, sqrd-1, phy-2* |
| Response to stress | 66 | 15 | 8.37E-11 | *hsp-70, cyp-37B1, cdr-2, nhr-115, ctl-1, mtl-1, ugt-44, nhr-57* |
| Transition metal ion binding | 62 | 14 | 1.07E-10 | *nhr-228, cyp-37B1, nurf-1, egl-9, mtl-1* |
| Small molecule metabolic process | 43 | 10 | 2.64E-07 | *cysl-3, pck-1, ugt-17, mmaa-1, mai-1, cysl-2* |
| Cellular response to unfolded protein | 13 | 3 | 1.21E-06 | *hsp-16.1, xbp-1, pek-1* |

The enriched biological terms for genes down-regulated by short-term hypoxia were single-organism metabolic process, monocarboxylic acid metabolic process, and oxidation-reduction process (Table 2). The representative genes for these functional groups were listed in Table 2. The full lists of genes for these functional groups were provided in S5 Table.

**Table 2.**
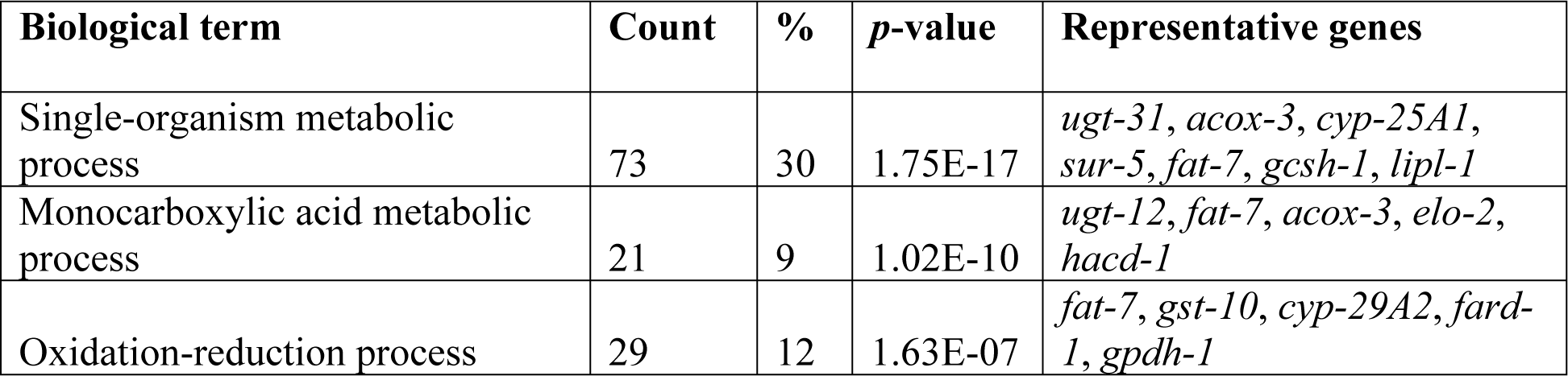
Enriched biological terms for genes down-regulated by short-term hypoxia in N2.

To further validate the data, we referred to genes that had been shown to be hypoxia-responsive in other published studies. RNase protection, RNA blot and real-time qRT-PCR assays had shown that F22B5.4, *nhr-57*, *fmo-12*/*fmo-2*, *egl-9*, *phy-2*, *cah-4*, K10H10.2/*cysl-2*, F26A3.4, C12C8.1/*hsp-70*, *acs-2* and *pck-1* were induced by hypoxia in N2 [17–20]. As expected, the microarray analyses described herein showed that all of these genes were induced by 2 hours of hypoxia in N2 (S2 Table). We also compared this dataset to a prior microarray study that identified 490 genes as hypoxia responsive when L3-stage N2 worms were treated with 4 hours of 0.1% oxygens [18]. While the larval stage and duration of hypoxia treatment were different, we were able to identify 50 genes that exhibited hypoxia-dependent changes in gene expression in these two experiments. The overlap was significant (*p*-value = 2.46E-10, by Fisher’s exact test), suggesting that both experiments identified gene expression changes responding to moderate hypoxia. In sum, the consistency with prior studies supported that our microarray data and analyses were promising.

### HIF-1-dependent short-term hypoxia responses

We next asked which short-term hypoxia-responsive gene expression changes were dependent upon HIF-1. To do this, we compared the hypoxia responses in N2 and *hif-1(ia04)* loss-of-function mutants, and this identified 124 genes whose hypoxia responses were different in *hif-1(ia04)* relative to N2 (S6 and S7 Tables). Among these, 64 genes were positively regulated by HIF-1 under hypoxia (their hypoxia inductions were higher in N2 than in *hif-1(ia04)*) (S6 Table), and 60 genes were negatively regulated by HIF-1 under hypoxia (their hypoxia inductions were lower in N2 than in *hif-1(ia04)*) (S7 Table). The heat maps in Fig 1 illustrate their hypoxia inductions in N2 and *hif-1(ia04)*, and the relative inductions (N2/*hif-1(ia04)*).

**Fig 1.**
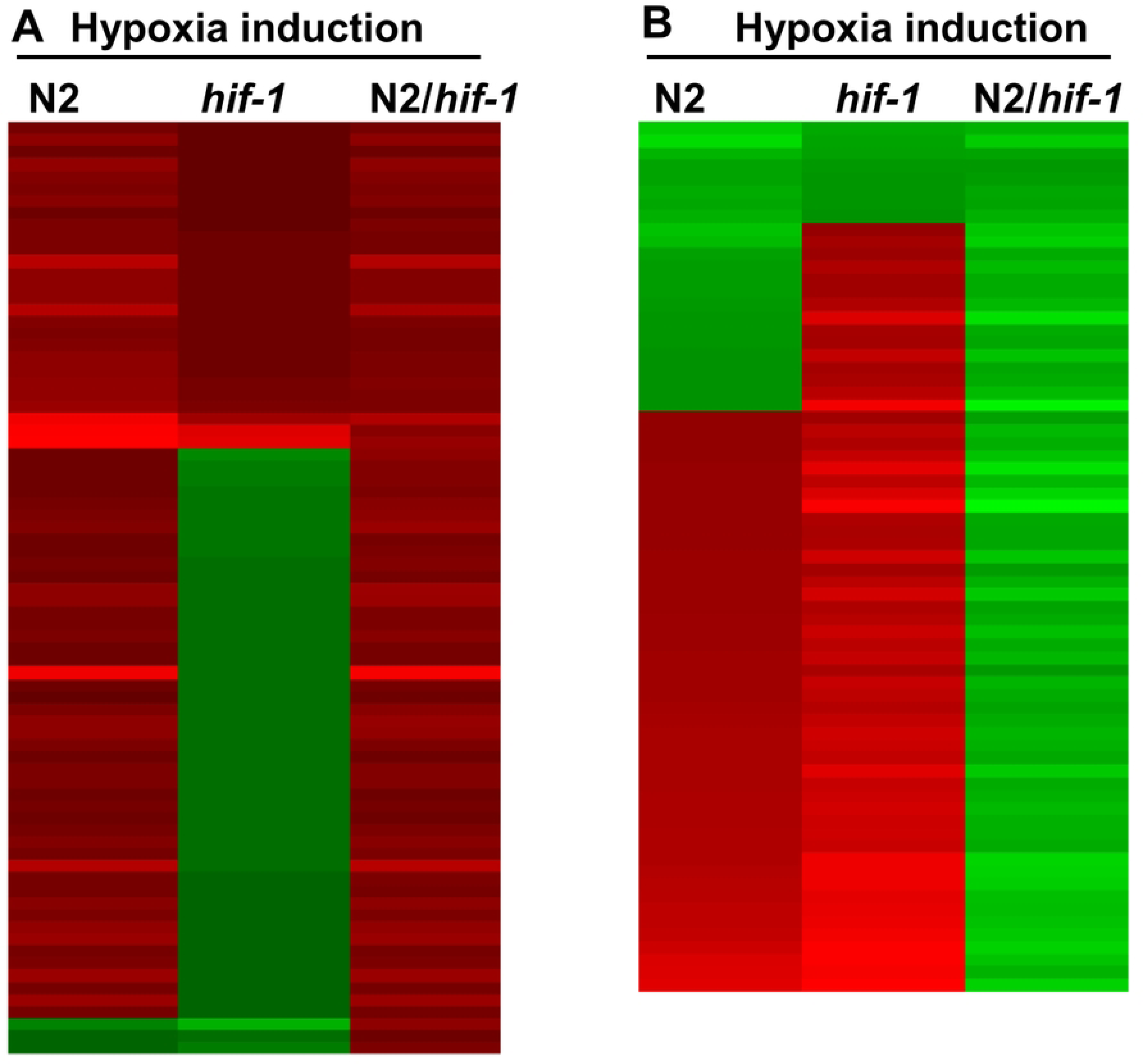
Hypoxia inductions of genes regulated by HIF-1 under hypoxia. (A, B) The heat map illustrations of hypoxia-dependent changes in gene expression for which HIF-1 was a positive regulator (A) or a negative regulator (B). Values < 0 were green, values > 0 were red. The color intensities corresponded to the induction levels in S6 (for Fig 1A) and S7 (for Fig 1B) Tables.

The enriched biological functions for genes positively regulated by HIF-1 under hypoxia were oxidation-reduction process, metabolic pathways, alpha-amino acid metabolic process, fatty acid metabolic process, and sulfur metabolism (Table 3). Several genes with important functions drew our attention. For example, *cysl-2*, *ethe-1* and *sqrd-*1 had been shown to have roles in detoxifying hydrogen sulfide (H_2_S) and hydrogen cyanide (HCN) [21], and *sqrd-*1 also maintained translation in H_2_S [22]. *mce-1*, *mmcm-1*, ZK550.6, *acox-1.6, gbh-2* and *fat-5* had been shown to be involved in lipid metabolism, and *asns-2*, *cysl-*2 and *ddo-1* had central roles in amino acid metabolism. *pck-*1 was integral to gluconeogenesis [20, 23], and *fmo-1* and *fmo-2* were involved in phase I detoxification. The *phy-2* (prolyl 4-hydroxylase) had been shown to have roles in cuticle collagen synthesis [24]. These findings were well aligned with previously published real-time qRT-PCR experiments that had shown that *pck-1*, *cysl-2* and *phy-2* were positively-regulated by HIF-1 under hypoxia [20].

**Table 3.**
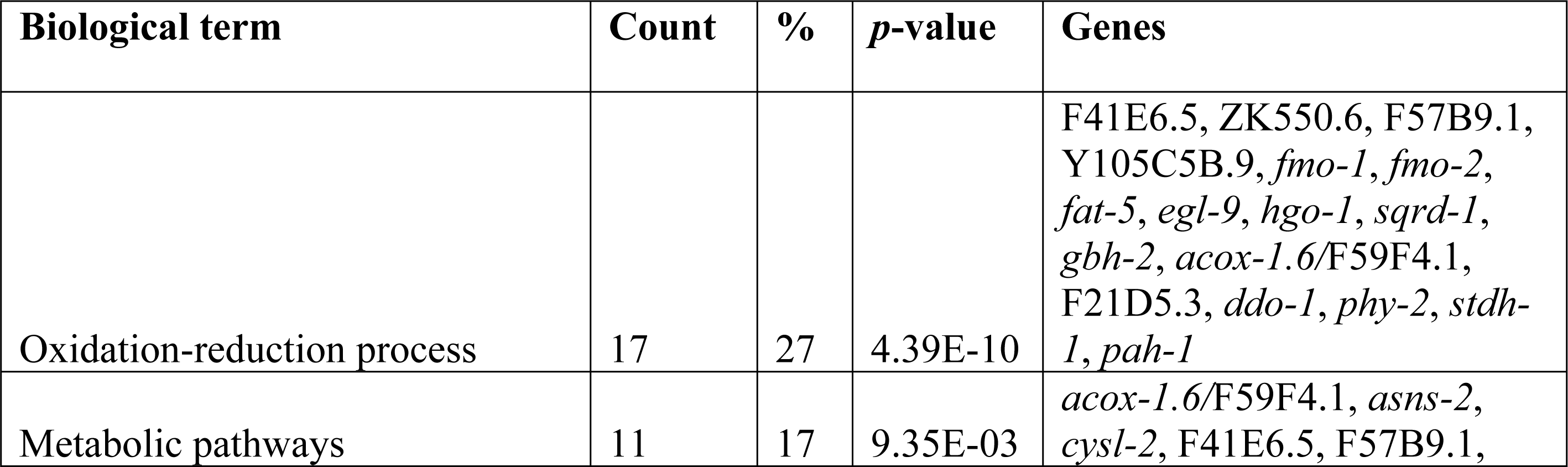
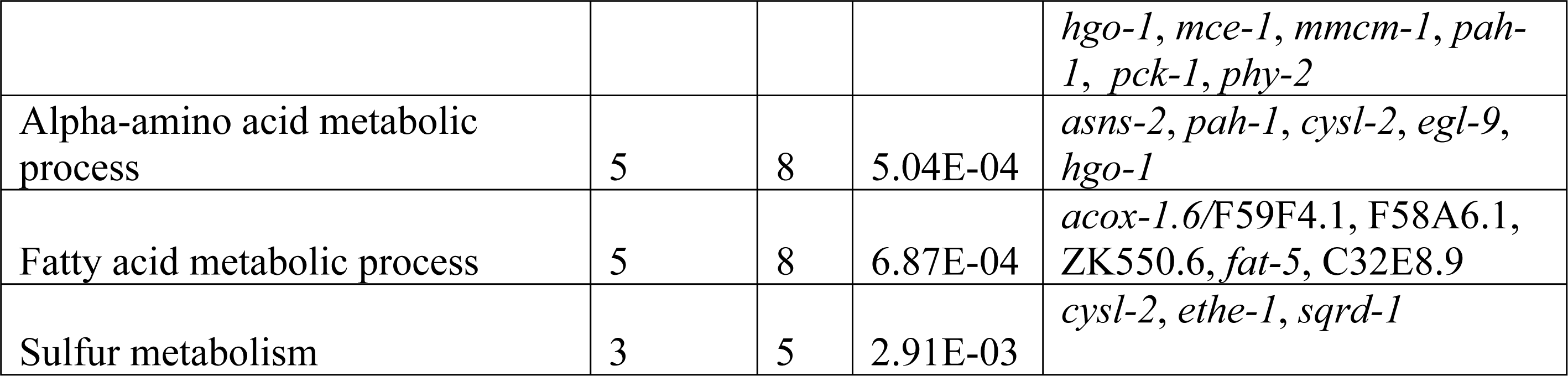
Enriched biological terms for genes positively regulated by HIF-1 under short-term hypoxia.

| Biological term | Count | % | p-value | Genes |
| --- | --- | --- | --- | --- |
| Oxidation-reduction process | 17 | 27 | 4.39E-10 | F41E6.5, ZK550.6, F57B9.1, Y105C5B.9, *fmo-1*, *fmo-2*, *fat-5*, *egl-9*, *hgo-1*, *sqrd-1*, *gbh-2*, *acox-1.6*/F59F4.1, F21D5.3, *ddo-1*, *phy-2*, *stdh-1*, *pah-1* |
| Metabolic pathways | 11 | 17 | 9.35E-03 | *acox-1.6* /F59F4.1, *asns-2*, *cysl-2*, F41E6.5, F57B9.1, *hgo-1, mce-1, mmcm-1, pah-1, pck-1, phy-2* |
| Alpha-amino acid metabolic process | 5 | 8 | 5.04E-04 | *asns-2, pah-1, cysl-2, egl-9, hgo-1* |
| Fatty acid metabolic process | 5 | 8 | 6.87E-04 | *acox-1.6/F59F4.1, F58A6.1, ZK550.6, fat-5, C32E8.9* |
| Sulfur metabolism | 3 | 5 | 2.91E-03 | *cysl-2, ethe-1, sqrd-1* |

The enriched biological functions for genes negatively regulated by HIF-1 under hypoxia were response to stress, and organic acid metabolic process (Table 4). Genes in the response to stress group included F-box A protein genes (*fbxc-58* and *fbxc-60*), infection response gene (*irg-2*), TIR domain protein gene *tir-1*, and UDP-glucuronosyl transferase gene *ugt-44*, and others. Genes in the organic acid metabolic process group included genes for lipid metabolism (*acdh-2*, *acs-2*, *elo-2*, *hacd-1* and *sur-5*), and others (Table 4).

**Table 4.** Enriched biological terms for genes negatively regulated by HIF-1 under short-term hypoxia.

| Biological term | Count | % | p-value | Genes |
| --- | --- | --- | --- | --- |
| Response to stress | 22 | 37 | 1.42E-10 | C49G7.10, *ckb-2, elo-2, F13A7.11, F18H3.4, F41B4.3, F56D2.5, fbxa-60, fbxc-58, ipla-2, irg-2, K09D9.1, nipi-3, parg-2, skr-3, T24C4.4, tir-1, tmem-135, ugt-44, warf-1, Y41C4A.11, Y43F8B.2* |
| Organic acid metabolic process | 9 | 15 | 1.25E-04 | *acdh-2, acs-2, ddo-2, elo-2, hacd-1, sur-5, ugt-25, ugt-28, ugt-44* |

### Genes responsive to both short-term and persistent HIF-1 activities

We anticipated that some of the genes that were responsive to short-term hypoxia would also be differentially expressed in mutants that over-expressed HIF-1 targets. To explore this question further, we compared the findings summarized in S2 and S3 Tables (genes up-regulated or down-regulated by 2-hour hypoxia) with genes that were mis-regulated in the four HIF-1 negative regulator mutants (*vhl-1(ok161)*, *rhy-1(ok1402)*, *egl-9(sa307)*, and *swan-1(ok267);vhl-1(ok161)* double mutants (See the related study entitled “Transcriptome analyses describe the consequences of persistent HIF-1 activation and reveal complementary roles for HIF-1 and DAF-16 in hypoxia survival in *Caenorhabditis elegans*”). This led to the identification of 23 genes were both positively regulated by HIF-1 under short-term hypoxia and up-regulated in all the four HIF-1 negative regulator mutants, in addition to 3 genes were both negatively regulated by HIF-1 under short-term hypoxia and down-regulated in all the four mutants (Tables 5 and 6). The molecular functions of the 23 genes positively regulated by both short-term and persistent HIF-1 activities were diverse, including genes for lipid metabolism (*mce-1*, *mmcm-1*, ZK550.6 and *gbh-2*), H_2_S and HCN detoxification (*cysl-2*, *ethe-1* and *sqrd-*1), gluconeogenesis (*pck-1*), and protein synthesis regulation (*efk-1*), as well as collagen synthesis (*phy-2*), and others. The 3 genes negatively regulated by both short-term and persistent HIF-1 activities were T28A11.2 (hypothetical protein), *acdh-2* (Acyl CoA dehydrogenase), and *acs-2* (fatty acid CoA synthetase family).

**Table 5.**
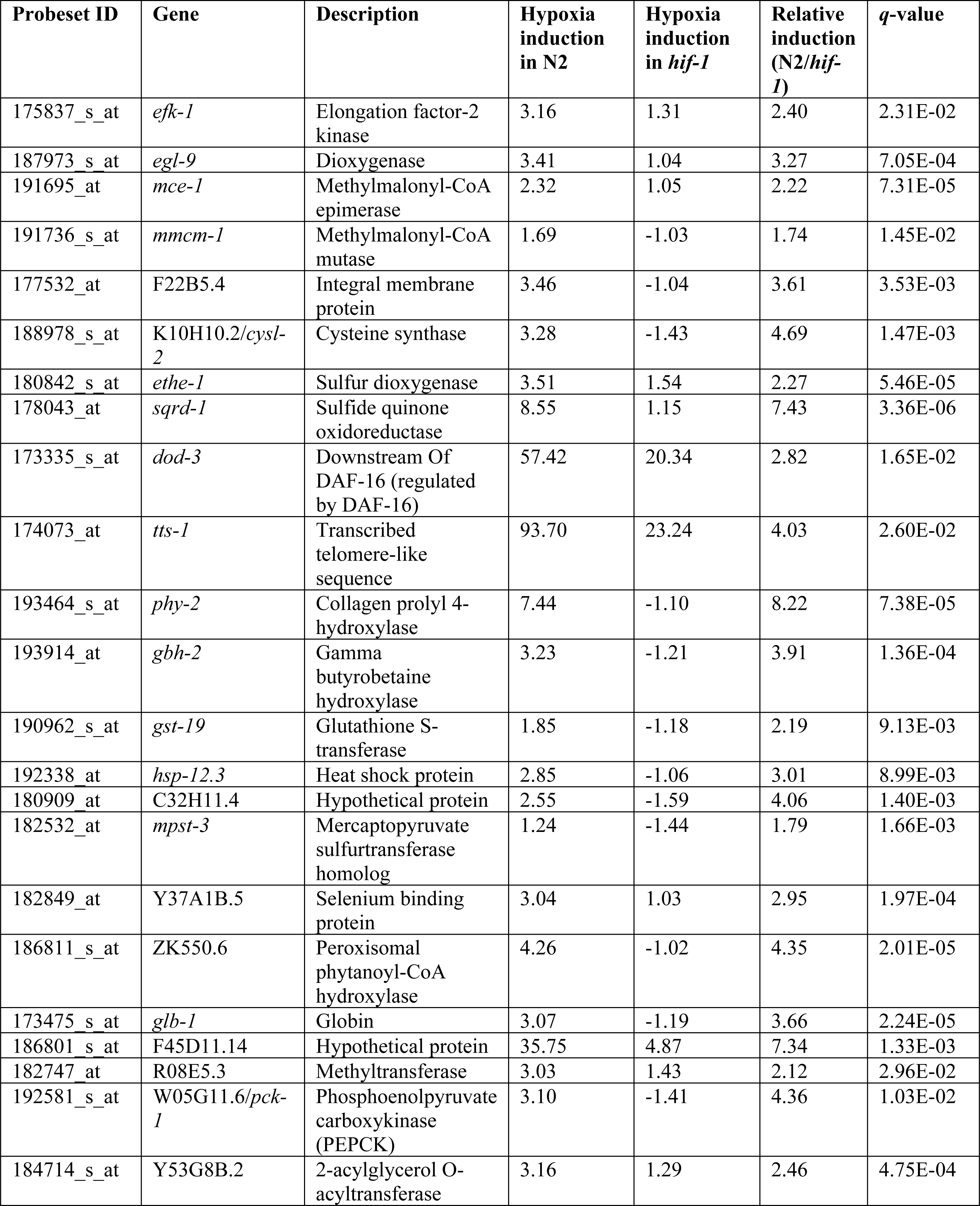
Hypoxia responses of genes positively regulated by both short-term and persistent HIF-1 activities.

**Table 6.** Hypoxia responses of genes negatively regulated by both short-term and persistent HIF-1 activities.

| Probeset ID | Gene | Description | Hypoxia-induction in N2 | Hypoxia induction in *hif-1* | Relative induction (N2/*hif-1*) | *q*-value |
| --- | --- | --- | --- | --- | --- | --- |
| 184849_at | T28A11.2 | Hypothetical protein | -1.28 | 1.47 | -1.88 | 1.28E-02 |
| 173996_at | *acdh-2* | Acyl CoA dehydrogenase | 1.05 | 3.22 | -3.08 | 1.17E-02 |
| 174675_at | *acs-2* | Fatty acid CoA synthetase family | 5.29 | 16.74 | -3.17 | 1.07E-02 |

### Effects of HIF-1-dependent hypoxia-responsive genes on hypoxia adaptation

To further understand HIF-1 function for hypoxia adaptation, we examined the requirement for genes that were induced by hypoxia in a HIF-1-dependent manner, with a focus on genes that had not been tested for their effects on hypoxia survival or had not been shown to have essential roles in development in room air. The actual number of genes tested was further limited by the availability of the mutant and RNAi strains. Twenty-seven mutants or RNAi treatments were examined to test 23 genes, and these genes had functions in multiple biological processes, including lipid metabolism, protein and amino acid metabolism, detoxification and stress response, ion transport, oxygen binding, vitamin biosynthesis, cellular signaling, protein translation regulation and collagen synthesis.

To assay the effects of a particular gene on hypoxia development and survival, we compared the abilities of animals to survive embryogenesis and larval development in hypoxia *versus* room air. These data are illustrated in Figs 2 and 3 (see also S8 Table). Wild-type N2 or N2;control RNAi (feeding L4440 empty vector) and *hif-1(ia04)* mutants were used as controls. As expected, N2 and N2;control RNAi animals were tolerant to hypoxia: their survival rates did not decrease under hypoxia compared to room air (*p*-values > 0.05). By contrast, *hif-1(ia04)* mutants were sensitive to hypoxia: only 78% hatched and 18% survived to adulthood in hypoxic conditions (\*\**p*-values < 0.01) (Fig 2, Fig 3 and S8 Table).

**Fig 2.**
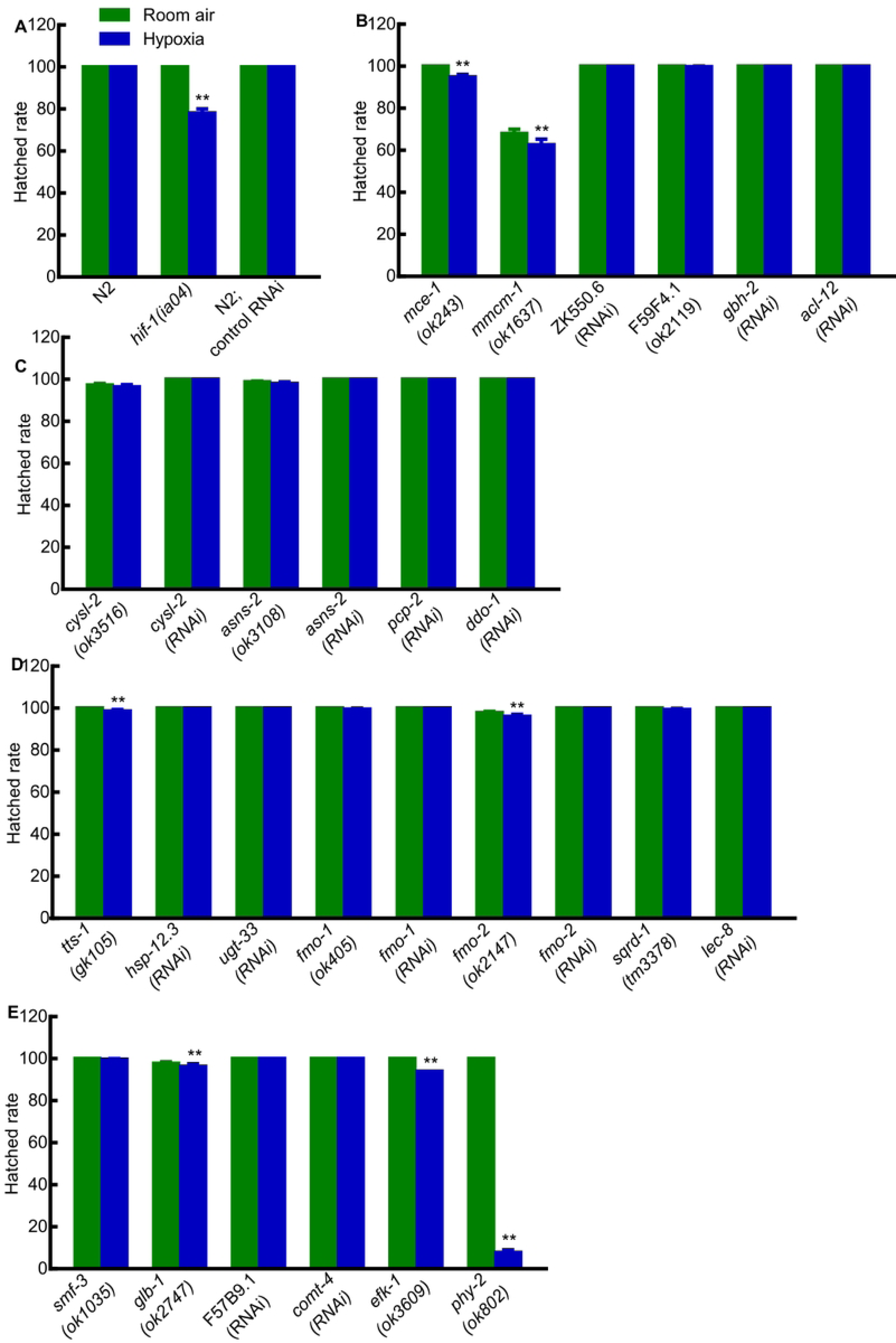
Effects of HIF-1-dependent hypoxia-responsive genes on embryogenesis. (A) Hatched rates of N2, *hif-1(ia04)* and N2;control RNAi in room air and hypoxia. (B-E) Hatched rates in room air and hypoxia for animals lacking specific genes related to (B) lipid metabolism; (C) amino acid metabolism; (D) detoxification and stress response; (E) ion transport (*smf-3*), oxygen binding (*glb-*1), vitamin biosynthesis (F57B9.1), cellular signaling (*comt-4*), protein translation regulation (*efk-1*) and collagen synthesis (*phy-2*). Values were mean ± SEM calculated from three biological replicates. The total number of animals assayed from three biological replicates for each strain in room air or hypoxia ranged from 205 to 661. \*\**p* < 0.01.

**Fig 3.**
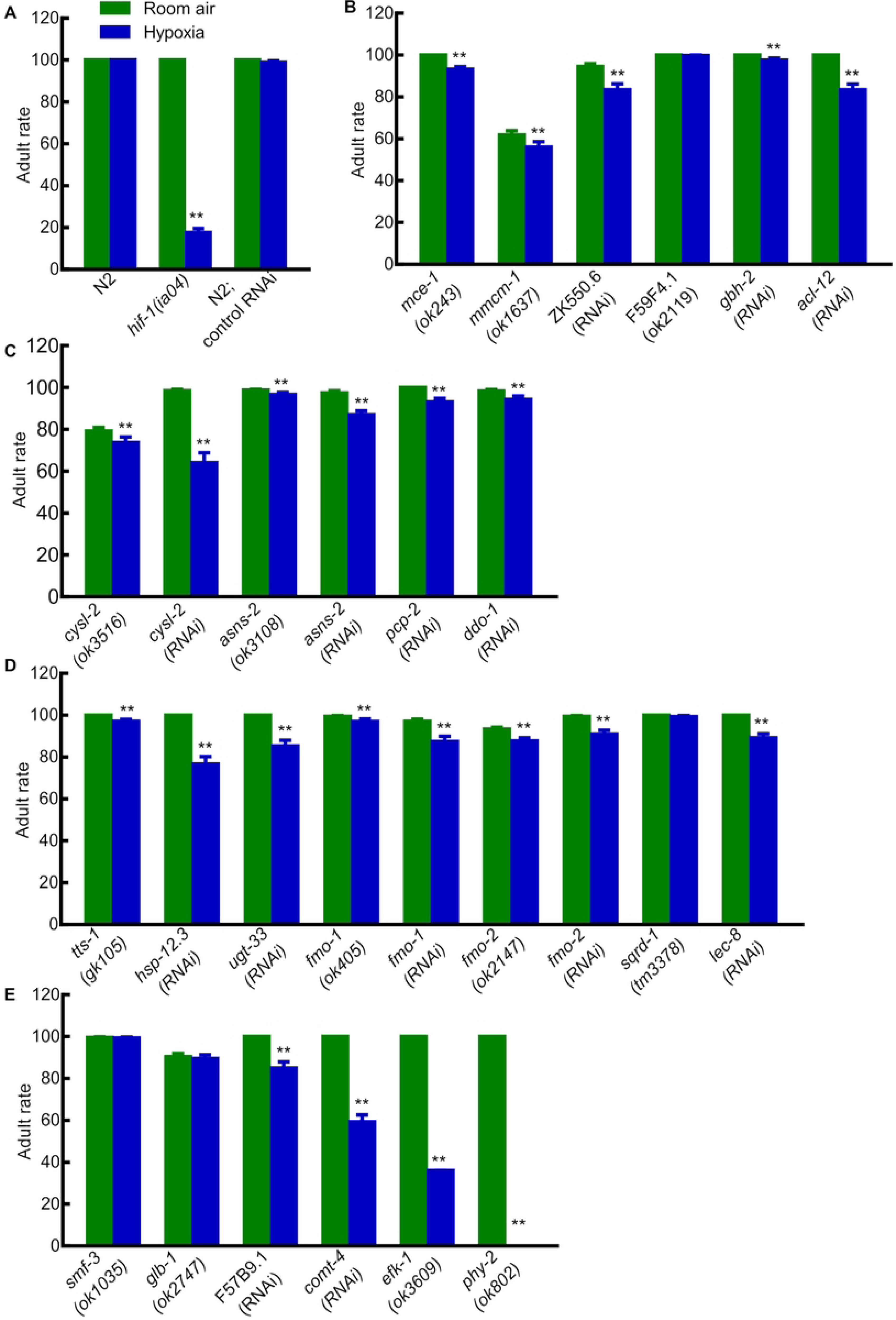
Effects of HIF-1-dependent hypoxia-responsive genes on survival to adulthood. (A) Rates of survival to adulthood for N2, *hif-1(ia04)* and N2;control RNAi in room air and hypoxia. (B-E) Rates of survival to adulthood in room air and hypoxia for animals lacking specific genes related to (B) lipid metabolism; (C) amino acid metabolism; (D) detoxification and stress response; (E) ion transport (*smf-3*), oxygen binding (*glb-*1), vitamin biosynthesis (F57B9.1), cellular signaling (*comt-4*), protein translation regulation (*efk-1*) and collagen synthesis (*phy-2*). Values were mean ± SEM calculated from three biological replicates. The total number of animals assayed from three biological replicates for each strain in room air or hypoxia ranged from 205 to 661. \*\**p* < 0.01.

Among the 27 mutants or RNAi conditions tested, 7 showed decreases in embryonic viability under hypoxia compared to room air, and 23 showed decreases in larval viability (\*\**p*-values < 0.01). The hypoxia effect of HIF-1-depent hypoxia-responsive genes on larval viability was stronger than that on hatched rate. Some mutant animals could hatch under hypoxia, but could not make to adulthood. In agreement with prior studies [17, 18], most of the genes tested had lesser roles in hypoxia survival than *hif-1* (hypoxia treatment decreased their embryonic or larval viability by no more than 20%). Notably, mutations in *hsp-12.3*, *cysl-2*, *comt-4*, *efk-1* or *phy-2* strongly impacted hypoxia survival, reducing survival to adulthood 23%, 34%, 41%, 64% and 100%, respectively, under hypoxia compared to room air.

### Identifying the direct targets of HIF-1 by ChIP-seq

To identify the genome sequences bound by HIF-1, we performed co-immunoprecipitation experiments. While there are six splicing isoforms of *hif-1*, the isoform a (*hif-1a*) had been shown to be essential for longevity and stress resistance [23, 25, 26]. Accordingly, we identified DNA sequences that co-immunoprecipitated with an epitope-tagged version of this HIF-1 isoform. From both biological replicates, we identified 94 reproducible HIF-1 binding peaks (FDR ≤ 0.05 and fold enrichment ≥ 1.6). The summaries of these peaks (including peak coordinates, sizes, distributions, ChIP and input tag counts, fold enrichments, target genes and their expressions under hypoxia and in the HIF-1 negative regulator mutants, et al.) are provided in S9 Table. Sequences co-immunoprecipitated with HIF-1 were provided in S1-S6 Files, organized by chromosomes and peak coordinates, one chromosome one file. ChIP signals were visually verified in IGB (Integrated Genome Browser) and provided in S1-S6 Figs, organized by chromosomes and target genes, one chromosome one file. As representatives, the IGB ChIP signals for *pqn-44* (prion-like-(Q/N-rich)-domain-bearing protein), *hsp-70* (heat shock protein), *nurf-*1 (nucleosome remodeling factor complex homolog), *efk-1* (eukaryotic elongation factor 2 kinase), *sqrd-1* (sulfide quinone oxidoreductase) and F19B2.5 (SNF2_N domain-containing protein) were presented in Fig 4. These six genes were among the HIF-1 direct targets whose expressions were changed by hypoxia or in the HIF-1 negative regulator mutants (see below Table 8). HIF-1 bound at different locations relative to these genes. The binding regions for *pqn-44* and *nurf-1* were within the coding regions. The HIF-1 binding site near *hsp-70* was upstream of the transcription start sites and overlapped with 5’ UTRs. The HIF-1 binding sites for *efk-1*, *sqrd-1* and F19B2.5 were also upstream of the transcription start sites (Fig 4). Among the HIF-1 direct targets whose expressions were changed by hypoxia or in the HIF-1 negative regulator mutants, *efk-1* was of particular interest. As shown in Figs 2 and 3, animals lacking *efk-1* function are less able to survive hypoxia. Gene expression studies show that *efk-1* expression is regulated by hypoxia and HIF-1 (see below Table 8). We verified the HIF-1 binding region in the *efk-1* promoter by ChIP-qPCR. The enrichment of this region was 8-fold relative to the reference *sir-2* promoter region (Fig 5). Consistent with this finding, this region was also identified as a HIF-1 binding site by the ModERN project [27].

**Fig 4.**
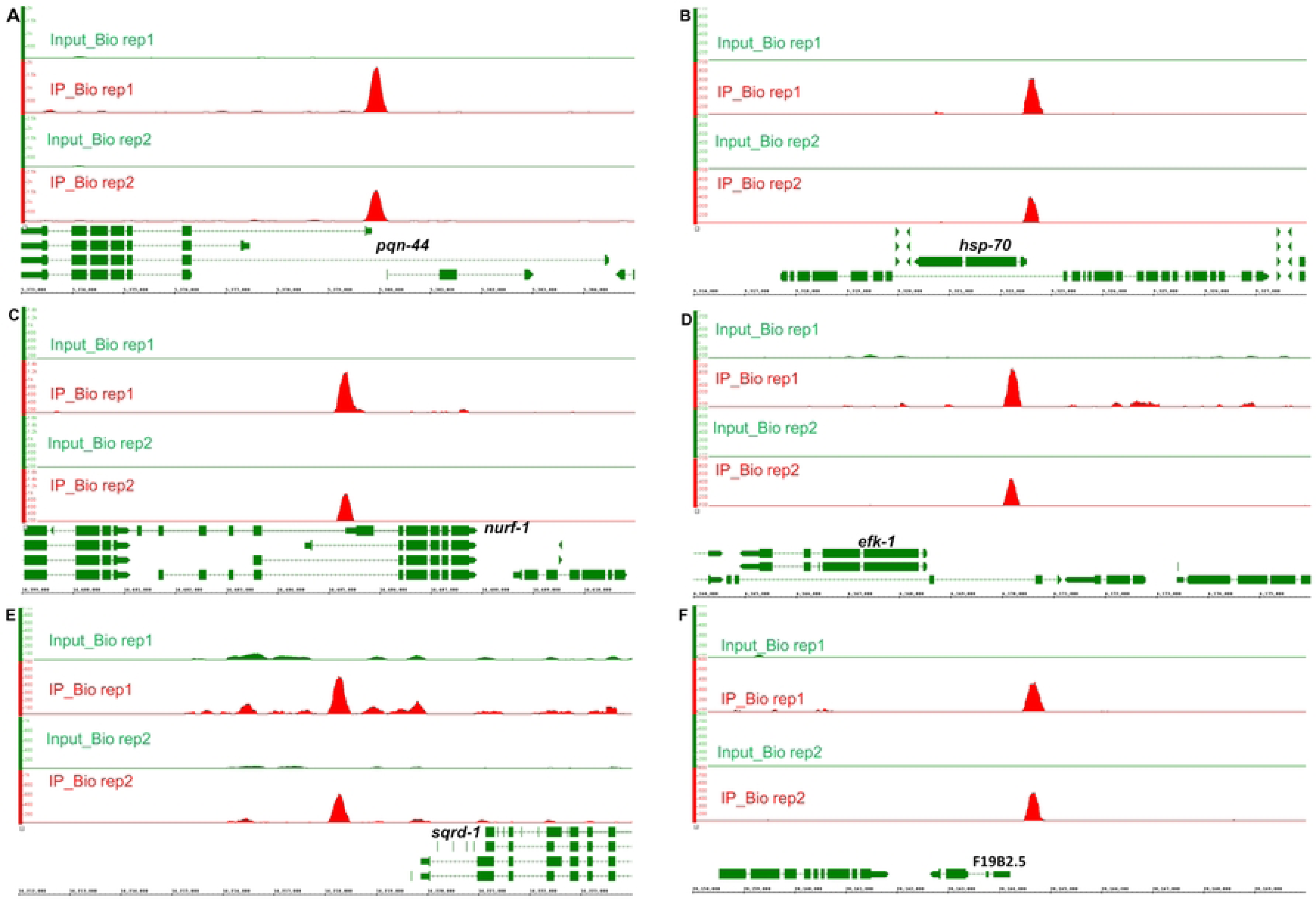
HIF-1 binding regions near six genes regulated by hypoxia or HIF-1. (A) A 600 bp HIF-1 binding region (chrI:5379600-5380199) within *pqn-44* (chrI:5372851-5384508). (B) A 400 bp HIF-1 binding region (chrI:9322400-9322799) upstream of *hsp-70* (chrI:9320325-9322519) and overlap with 5’ UTR. (C) A 600 bp HIF-1 binding region (chrII:14405000-14405599) within *nurf-1* (chrII:14390713-14407911). (D) A 400 bp HIF-1 binding region (chrIII:6170000-6170399) 1453 bp upstream of the transcription start of *efk-1* (chrIII:6164906-6168547). (E) A 600 bp HIF-1 binding region (chrIV:14218000-14218599) 1272 bp upstream of the transcription start of *sqrd-1* (chrIV:14219871-14224440). (F) A 600 bp HIF-1 binding region (chrV:20164400-20164999) 235 bp upstream of the transcription start of F19B2.5 (chrV:20162615-20164165). The images showed the IGB ChIP-seq signals in both biological replicates. For each binding region in each biological replicate, the minimum number for the y-axis scale was the normalized average input tag count, and the maximum number was the averaged ChIP tag count as provided in S9 Table.

**Fig 5.**
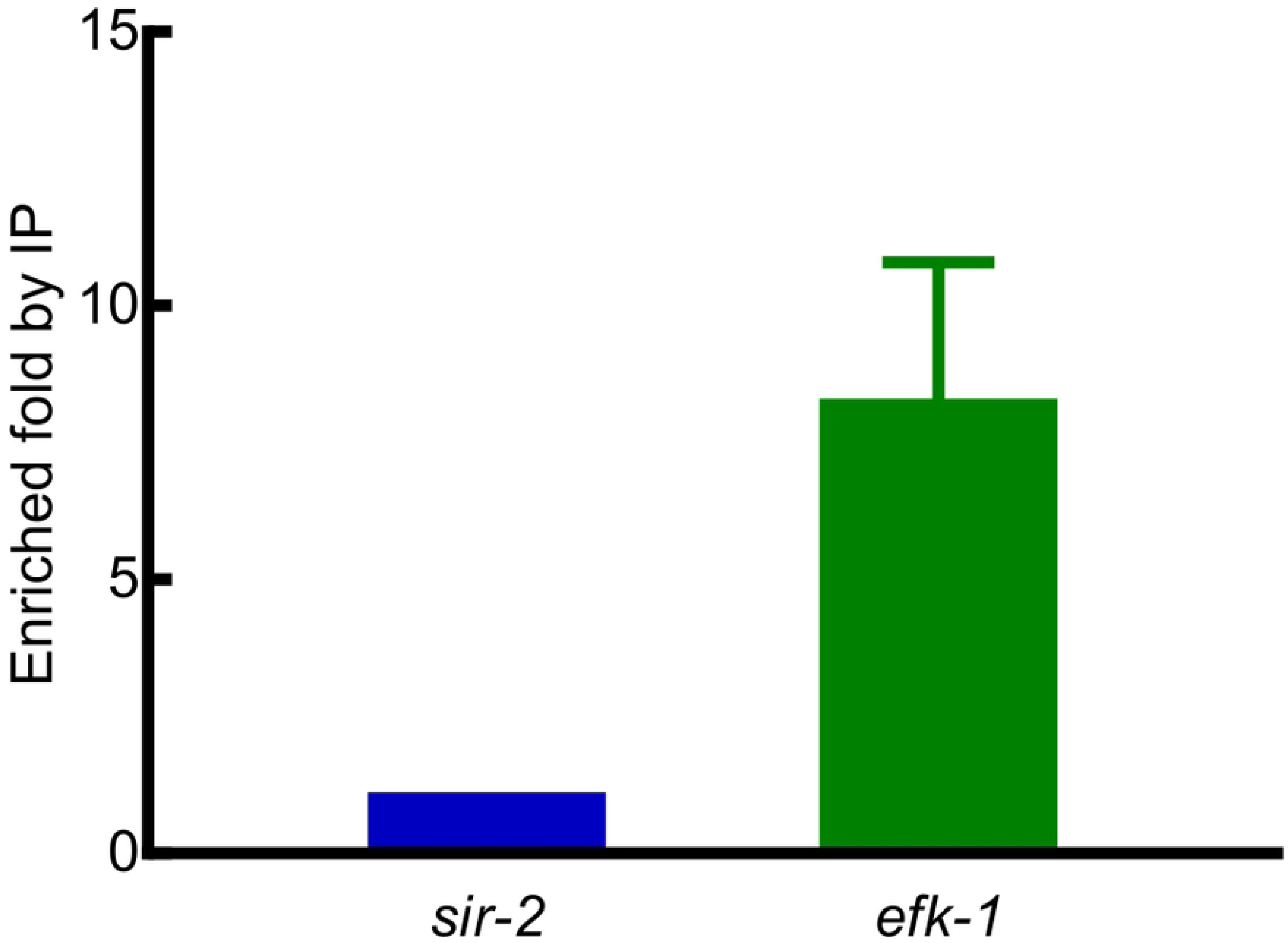
*efk-1* ChIP-qPCR. ChIP-qPCR to verify the HIF-1 binding region in the *efk-1* promoter. The *sir-2* promoter region was used as the reference. Values were mean ± SEM calculated from three biological replicates.

Most of the HIF-1 binding peaks were located in introns (34.38%), upstream of the transcription start sites (28.13%), or downstream of the transcription stop sites (23.96%). The rest of binding peaks lied upstream of the transcription start sites and overlap with 5’ UTRs (11.46%), or in the coding regions (1.04%), as well as in 3’ UTRs (1.04%) (Fig 6 and S9 Table). A majority of the HIF-1 binding peaks (60/94 = 64%) contained sequences similar to mammalian HRE (hypoxia response element) 5’-RCGTG-3’ (R = A or G), or the mandatory core HRE 5’-CGTG-3’ [28] (S9 Table and S1-S6 Files).

**Fig 6.**
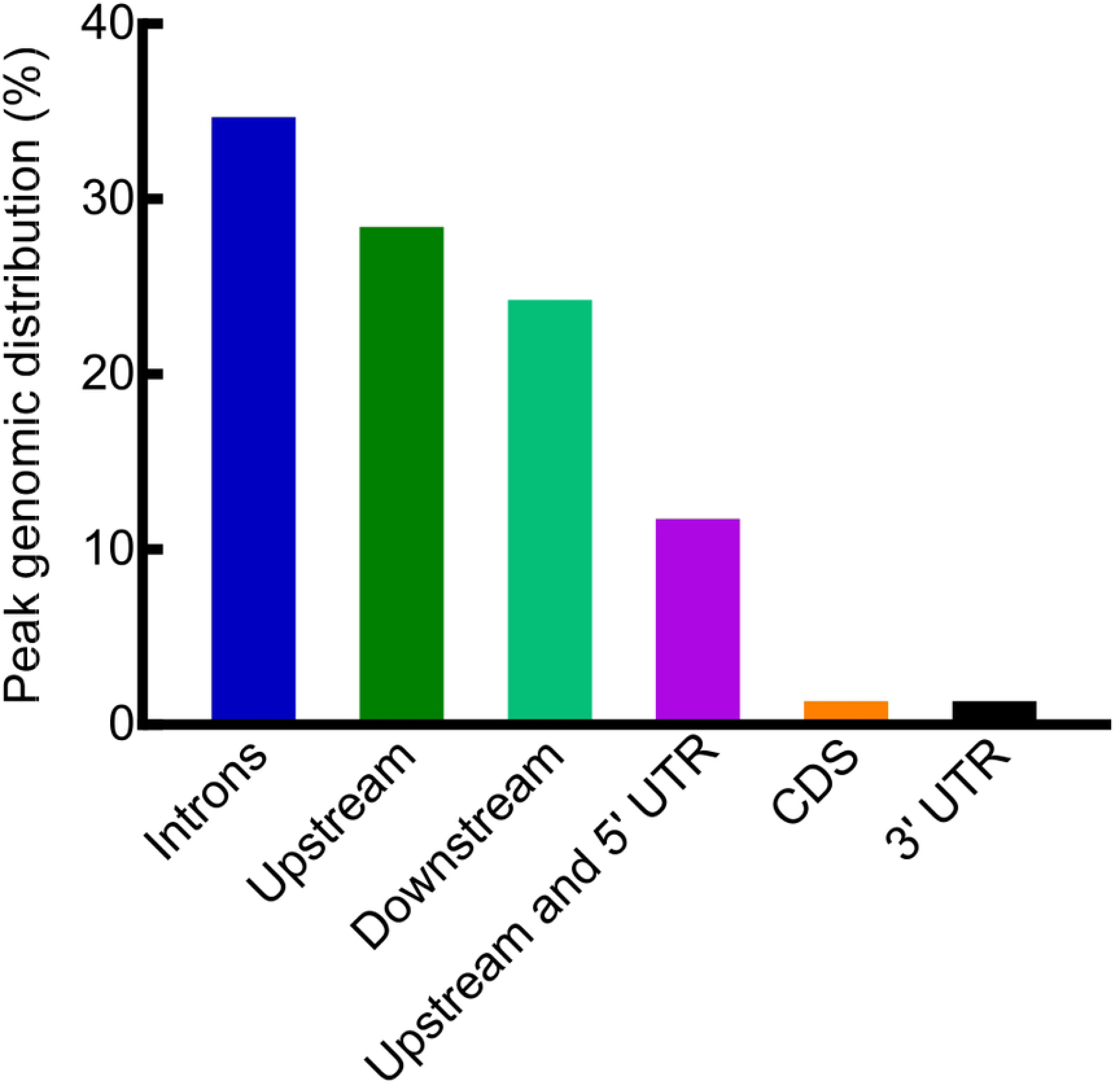
Genomic distributions of HIF-1 ChIP peaks. Genomic distributions of HIF-1 ChIP peaks relative to target genes. The detailed distributions for each peak were provided in S9 Table.

These 94 HIF-1 binding regions were annotated to 96 genes (S9 Table). In this study, these genes were treated as HIF-1 direct target genes. The enriched biological terms associated with HIF-1 direct target genes were protein processing in endoplasmic reticulum, response to stress, and cell recognition (Table 7). Genes in the protein processing in endoplasmic reticulum group included heat shock protein genes (*hsp-70*, *hsp-90* and *dnj-12*), and others. Genes in the response to stress group included heat shock protein genes (*hsp-16.2*, *hsp-16.41*, *hsp-70*, *hsp-90*, *hsp-110*, and *dnj-12*), *atg-*5 (autophagy protein 5), *tir-1* (TIR domain protein), and others. And genes in the cell recognition group included *cua-1* (CU (copper) ATPase), *madd-2* (muscle arm development defective), and others (Table 7).

**Table 7.** Enriched biological terms associated with HIF-1 direct target genes.

| Biological term | Count | % | p-value | Genes |
| --- | --- | --- | --- | --- |
| Protein processing in endoplasmic reticulum | 7 | 7.29 | 5.74E-06 | *dnj-12*, *ero-1*, F44E5.4, F44E5.5, *hsp-70*, *hsp-90*, *unc-23* |
| Response to stress | 14 | 14.58 | 4.36E-04 | *atg-5*, *cpr-3*, *daf-25*, *dnj-12*, *ero-1*, *hsp-16.2*, *hsp-16.41*, *hsp-70*, *hsp-90*, *hsp-110*, *irg-1*, *tir-1*, *tank-1*, Y56A3A.33 |
| Cell recognition | 4 | 4.17 | 5.85E-03 | *cua-1*, *madd-2*, Y105E8B.9, ZC262.7 |

**Table 8.** Genes that co-immunoprecipitated with HIF-1 and were shown to be regulated by HIF-1.

| Expression | Count | Genes |
| --- | --- | --- |
| Positively regulated by HIF-1 under short-term hypoxia | 3 | *efk-1*, *sqrd-1*, W03F9.1 |
| Negatively regulated by HIF-1 under short-term hypoxia | 1 | *tir-1* |
| Up-regulated in at least one mutant that stabilizes HIF-1 | 14 | Y105E8B.9, F44E5.5, F44E5.4, *nurf-1*, *efk-1*, C46C2.5, *sqrd-1*, *hsp-16.41*, *hsp-16.2*, *irg-1*, *ari-1.4/tag-349*, F54D5.12, *nurf-1*, *cpr-3* |
| Down-regulated in at least one mutant that stabilizes HIF-1 | 9 | C50A2.3, C46C2.5, F54D5.12, *oat-1*, Y105C5B.5, F19B2.5, M04F3.3/*kin-35*, Y94H6A.5, C23H5.8 |

### Genomic regions that co-immunoprecipitate with HIF-1 map to genes that are differentially regulated by conditions or mutations that alter HIF-1 function

Among the 96 gene regions that co-immunoprecipitated with HIF-1, 24 mapped to genes that were regulated by HIF-1 under short-term hypoxia or in mutants with constitutively active HIF-1 (single mutants of *vhl-1(ok161)*, *rhy-1(ok1402)* and *egl-9(sa307)*, and *swan-1(ok267);vhl-1(ok161)* double mutants) (Table 8).

An interesting feature emerging from the expression patterns of HIF-1 direct targets was that they responded to HIF-1 in differing contexts. Some genes responded to HIF-1 under short-term hypoxia (for example, W03F9.1 and *tir-*1), while others were differentially expressed in mutants in which HIF-1 was constitutively active (for example, *oat-1* and *cpr-3*). As noted in Table 8, some of the genes that coimmuniprecipitaed with HIF-1 had been shown to respond to HIF-1 under both short-term hypoxia and in the mutants with constitutively active HIF-1 (like *efk-1* and *sqrd-1*).

## Discussion

Transcription factors HIFs are the master regulators of hypoxia response. Identification of direct HIF-1 targets is a major step towards more fully understanding the transcriptional networks controlled by *C. elegans* HIF-1. Here, we add to the literature that describes hypoxia-responsive gene expression, in ways that provide new insights to HIF-1 mediated immediate hypoxia response. By cross-referencing the genes that are differentially regulated by hypoxia or HIF-1 with those genomic regions that co-immunoprecipitate with HIF-1, we can describe the direct and downstream targets of HIF-1 with greater confidence.

### Refined understanding of HIF-1 mediated short-term hypoxia response

The transcriptome analyses confirmed and expanded prior studies that have investigated the roles of HIF-1 in hypoxia response. The results of each study is, of course, influenced by the hypoxia regimen, the larval stage tested, and the genetic background. Here, we opted to use multiple strategies to induce HIF-1 through hypoxia (0.5% O2 for 2 hours) or through loss-of-function mutations in HIF-1 negative regulators. By compiling and cross-referencing these data sets, we can develop a list of high-confidence HIF-1 targets. As a complementary experimental strategy, we identified genomic regions that co-immunoprecipitated with HIF-1.

These data sets provide insights to the myriad of hypoxia-responsive functions regulated by HIF-1. Some of these functions are evolutionarily conserved, especially stress response and energy metabolism. There are notable overlaps between genes regulated by HIF-1 under hypoxia with those regulated by DAF-16 [29] (*p*-value = 6.80E-10, by Fisher’s exact test) or with those regulated by H_2_S [30] (*p*-value = 4.04E-05, by Fisher’s exact test), and this suggests that HIF-1 has broad roles in stress response. In the wild, *C. elegans* can live in hypoxic microenvironments, and regulations of stress response pathways are likely to be critical to intergenerational survival [31]. Some of the key findings are summarized in Table 8, and multiple lines of evidence point to key direct targets of HIF-1. These include elongation factor kinase *efk-1*, the heat shock factor *hsp-16.2*, the mitochondrial sulfide quinone oxidoreductase *sqrd-1* and the nucleosome remodeling factor *nurf-1*.

While this provides an important foundation upon which to build, the number of direct HIF-1 targets are no doubt much higher than the 96 genes described in S9 Table. Any one binding site will only be occupied some fraction of time. Another limitation is that, in lieu of a reliable HIF-1-specific antibody, we used an epitope-tagged version of HIF-1. While there are good reasons for focusing on the a isoform, it is likely that other isoforms have biological functions as well. With time and improved technologies, additional studies will add to this core dataset.

## Materials and methods

### Strains

The wild-type *C. elegans* used in this study was N2 Bristol. The mutant strains used in this study were listed in S10 Table. All the worms were maintained at 21°C using the standard methods [32].

### Gene expression microarray experiment

Randomized complete block design was followed for the microarray experiment, with three biological replicates treated as three blocks. Each block included eight treatments: N2 wild type, N2 wild type with hypoxia treatment, *hif-1*(*ia04)* loss-of-function mutants*, hif-1*(*ia04)* loss-of-function mutants with hypoxia treatment, *vhl-1(ok161)* loss-of-function mutants*, rhy-1(ok1402)* loss-of-function mutants, *egl-9(sa307)* loss-of-function mutants and *swan-1(ok267);vhl-1(ok161)* loss-of-function double mutants. For each treatment, about 1,000 synchronized L4-stage larvae were pooled as one experimental unit to get sufficient RNA for hybridization. Total RNA isolation was performed using Trizol (Invitrogen) and RNeasy Mini Kit (Qiagen). RNA quality was checked with an Agilent 2100 BioAnalyzer (Agilent Technologies). The RNA integrity numbers (RINs) for all the samples used in this study were greater than 9.0. The total RNA isolated from one experimental unit was hybridized onto one Affymetrix GeneChip® C. elegans Genome array (Affymetrix, part number 900383). Probe synthesis, labeling, hybridization, washing, staining and scanning were performed by the GeneChip facility at Iowa State University. In brief, the total RNA was synthesized to biotin-labeled aRNA using the GeneChip® 3’ IVT Express Kit (Affymetrix, part number 901229) and hybridized to the array.

The arrays were washed and stained in the GeneChip® fludics station 450 and scanned with GeneChip® scanner 3000 7G. The Affymetrix® GeneChip® Command Console™ (AGCC) software was used to generate probe cell intensity data (.CEL) files. The resulting CEL files were normalized and summarized using the robust multichip average (RMA) algorithm [33] in R package (R Core Team, Vienna, Austria, 2016). An analysis of variance (ANOVA) model was then fitted to the summarized expression measures, with the block (three levels) and the treatment (eight levels) treated as fixed effect factors following the experimental design.

Residual model diagnostics identified no severe violations of the model assumptions. Linear contrasts of treatment means were tested using the general F-test. To account for multiplicities of hypothesis testing, conservative estimates of false discovery rates (FDRs) were calculated according to the *q*-value procedure of Storey and Tibshirani [34]. Differentially expressed probesets were defined as *q*-value ≤ 0.05 and fold change ≥ 1.6. Probesets were converted to genes using the Affymetrix annotation file “Celegans.na36.annot.csv. To deal with redundancy and count the number of unique genes detected on the array, we kept one probeset per gene and one gene per probeset. In this way, the total number of unique genes detected on the array was 18, 011. For the purpose of reference, the original complete lists of gene(s) annotated to each probeset were kept in S1-S3, S6 and S7.

In this paper, we discuss HIF-1-dependent hypoxia responses. Gene expression changes in the mutants with constitutive HIF-1 activity will be described in a related study entitled “Transcriptome analyses describe the consequences of persistent HIF-1 over-activation in *Caenorhabditis elegans*”. The microarray raw and probeset summary data had been deposited to NCBI’s Gene Expression Omnibus, the accession number was GSE228851.

### Gene function annotation and enrichment analyses

DAVID tools (The Database for Annotation, Visualization and Integrated Discovery) [35, 36] (https://david.ncifcrf.gov) were used to annotate the enriched biological terms associated with microarray and ChIP-seq-selected genes. The enriched biological terms were at *p*-value < 0.01 with no correction.

### Heat maps

Heat maps for gene expression profiles were generated by the PermutMatrix graphical analysis program [37, 38]. Average linkage clustering was performed using the hypoxia induction values. Green color represented negative values, and red color represented positive values. The intensities of the colors represented the magnitudes of fold changes. Other parameters were set as default.

### Gene lists overlap testing

Fisher’s exact test was performed to test whether the overlap between two gene lists was significant or not. The total number of 18, 011 genes detected on the microarray was used as the population size. The significant overlap is at *p*-value < 0.001.

### HIF-1 chromatin immunoprecipitation sequencing (ChIP-seq)

The ChIP experiments were performed in the *egl-9(sa307)* loss-of-function mutant background to maintain HIF-1 stability and activity in room air. The strain used for the ChIP experiments was ZG434 (*egl-9(sa307);iaIS28[Phif-1::hif-1a::Myc::HA];hif-1(ia04)*). To use the commercially available ChIP grade anti-HA antibody (Abcam, cat. no. ab9110), an HA-tagged *hif-1a* transgene *iaIS28[Phif-1::hif-1a::Myc::HA]*) [25] was introduced into *egl-9(sa307)*, and the endogenous *hif-1* gene was knocked out. Detailed ChIP protocol was provided in S7 file. Briefly, synchronized L4-stage worms for each biological replicate were harvested in separate batches at separate times. Harvesting enough synchronized worms for HIF-1 ChIP experiment was laborious due to the egg-laying defect inherent to the *egl-9(sa307)* loss-of-function mutation. For each batch, about 10, 000 L4-stage worms were harvested and cross-linked in 2% formaldehyde at 21°C for 30 minutes. The ChIP-seq experiment was performed with two biological replicates. For each biological replicate, nuclear lysates from about 50,000 worms (pooled from 5 batches of worm collection) were sonicated using a Branson sonifer microtip to fragment the chromatin to 200-800 bp. Immunoprecipitated protein-DNA complexes were captured on protein A-Sepharose beads (Sigma) and eluted in elution buffer (1% SDS and 100 mM NaHCO3) at 65°C for 30 minutes. Following RNase treatment and cross-link reversal, the ChIP DNA was purified with the Qiagen MinElute Kit and stored at −20°C for sequencing in parallel with the corresponding input DNA. The ChIP-Seq library preparation and sequencing were performed by the Iowa State University DNA facility. In brief, NEXTflex™ ChIP-Seq Barcodes kit (Illumina compatible) (BIOO Scientific Corp., cat. no. 514123) was used to prepare multiplexed single-end genomic DNA libraries. The gel slices corresponding to the 200-300 bp maker were cut and purified. The purified DNA was amplified and sequenced in a single flow cell on the IlluminaHiSeq 2000 platform. The length of reads was 50 bp.

### HIF-1 ChIP-seq data analyses

The HIF-1 ChIP-seq raw and processed data had been deposited to NCBI’s Gene Expression Omnibus and the accession number was: GSE228846. The quality scores of the fastq reads for the input and ChIP DNA samples from both biological replicates were all above 30, indicating high quality sequencing data (S7 Fig). Reads were mapped to *C. elegans* reference genome ce11 using bowtie 2 with the default settings. Reads with mapping quality (MAPQ) score less than 10 or reads mapped to the mitochondrial genome were excluded. At the end, reads kept for peak calling were 19-43 million per sample. The kept reads were assigned to bins, the size of which was set at 200 bp to match the fragment length for Illumina sequencing. Bin-level read counts were analyzed by the R package MOSAiCS (MOdel-based one and two Sample Analysis and Inference for ChIP-Seq Data) to call peaks [39] (https://www.bioconductor.org/packages/release/bioc/manuals/mosaics/man/mosaics.pdf). The false discovery rate (FDR) was set at 0.05. Neighboring peaks were merged. The output peaks were further filtered with the following criteria: (1) minimum posterior probability ≤ 0.05; (2) averaged input tag count ≥ 10; (3) averaged ChIP tag count ≥ 10; and (4) fold enrichment (averaged ChIP tag count/normalized average input tag count) ≥ 1.6. The identified peaks were visually verified in IGB (Integrated Genome Browser) [40]. The WIG files for IGB were provided in GEO database (accession number GSE228846). Peaks identified by both biological replicates were treated as HIF-1 binding regions. Peaks were assigned to genes within 6 kb. Within this region, if there was a gene(s) differentially expressed under hypoxia or in the HIF-1 negative regulator mutants (*vhl-1(ok161)*, *egl-9(sa307)*, *rhy-1(ok1402)* and *swan-1(ok267);vhl-1(ok161)* mutants), the peak was assigned to this gene. We reasoned that a gene(s) showed expression change under these conditions was more possible to be a HIF-1 direct target than genes showed no expression changes. Otherwise the nearest gene was assigned to the peak. Most often (90 out of 96 genes), the assigned HIF-1 direct target was the nearest gene.

### ChIP-qPCR to verify the HIF-1 binding site in the *efk-1* promoter

The primers for ChIP-qPCR to verify the HIF-1 binding region in the *efk-1* promoter were the forward primer 5’-CAATCTGACCGAGCCGAATG-3’ and reverse primer 5’-AGGCCTTTCTCGATTTTCCA-3’. The amplicon was 172 bp and contained a HRE 5’-ACGTG-3’. The promoter region of *sir-2*, a gene not regulated by HIF-1 under short-term hypoxia or in the HIF-1 negative regulator mutants, was used as the reference. The primers for *sir-2* ChIP-qPCR were the forward primer 5’-AGATTGCTTCTTTGGCTGGA-3’ and reverse primer 5’-GTAACGCACCTTGCAACAGA-3’. The amplicon was 218 bp and did not contain HRE similar sequences. Three biological replicates were performed. qPCR quantification was performed using the efficiency-corrected comparative quantification method [41].

### Hypoxia development and survival assays

For each mutant genotype, the room air and hypoxia treatments were performed in parallel at 21°C. For each treatment, 20 young adults (one day after L4 molt) were used as parents to lay eggs on one NGM plate seeded with OP50 for 30 minutes. After counting the eggs laid, the plates were kept in room air or put into a sealed plexiglass chamber with constant hypoxic gas flow for 24 hours. Compressed air and 100% nitrogen were mixed to achieve 0.5% oxygen, and gas flow was controlled by an oxygen sensor [18]. After 24 hours, the un-hatched eggs were counted for both treatments. After that, the plates for both treatments were maintained in room air. The adult worms were counted 72 hours after the eggs had been laid. The data collection time points were set to match the development rate of N2 eggs in room air: they hatched within 24 hours and reached adulthood within 72 hours.

For RNAi strains, RNAi was induced by bacterial feeding as described [42, 43]. Except for F57B9.1, *gbh-2* and *comt-4*, the RNAi clones were purchased from the Ahringer RNAi library (Geneservice, Cambridge, UK) and validated by sequencing. The RNAi constructs for F57B9.1, *gbh-2* and *comt-4* were generated by cloning the coding regions into the L4440 double-T7 vector [42]. To generate the RNAi parent generation, 20 N2 adults maintained with OP50 were transferred to RNAi plates to lay eggs for 1 hour. Three days later, 20 young adults grown up from these eggs were randomly picked as RNAi parents to lay eggs on a new RNAi plate for 30 minutes for either room air or hypoxia treatment. The room air and hypoxia treatments were performed in parallel at 21°C. The downstream procedures for hypoxia treatment and counting the hatched/un-hatched eggs and adult/non adult animals were the same as those described above for the mutant strains.

The experiments were performed with three biological replicates. To test the effect of hypoxia on animal development and survival, the binary hatched *vs*. un-hatched or adult *vs.* non adult data were analyzed by fitting a generalized linear model using a logit link function with JMP 9 statistical software (SAS Institute Inc., Cary, NC, 2010). The replicate (three levels) and the treatment (two levels) were used as factors in the model. For situations in which such models were inappropriate, randomization tests were used.

## Acknowledgments

The experiments reported here were conducted as part of Dingxia Feng’s PhD thesis work, under the guidance of Dr. Jo Anne Powell-Coffman at Iowa State University. The work was supported by grant R01GM078424 from the National Institutes of Health to JP. Author DF conducted the experiments, performed the ChIP-seq data analyses and drafted the manuscript, and author LQ was instrumental to the statistical design and analyses of gene expression comparisons. The authors are grateful for mentors and thesis committee members who provided advice and guidance.

## Supporting information

**S1 Table. Gene expression for all the probesets under hypoxia and in the HIF-1 negative regulator mutants.**

**S2 Table. Genes up-regulated by short-term hypoxia in N2.**

**S3 Table. Genes down-regulated by short-term hypoxia in N2.**

**S4 Table. Enriched biological terms for genes up-regulated by short-term hypoxia in N2.**

**S5 Table. Enriched biological terms for genes down-regulated by short-term hypoxia in N2.**

**S6 Table. Genes positively regulated by HIF-1 under short-term hypoxia.**

**S7 Table. Genes negatively regulated by HIF-1 under short-term hypoxia.**

**S8 Table. Effects of HIF-1-dependent hypoxia responsive genes on hypoxia development and survival.**

**S9 Table. Direct targets for HIF-1 identified by ChIP-seq.**

**S10 Table. Descriptions of mutations used in this study.**

**S1 File. Sequences co-immunoprecipitated with HIF-1 on chromosome 1.**

**S2 File. Sequences co-immunoprecipitated with HIF-1 on chromosome 2.**

**S3 File. Sequences co-immunoprecipitated with HIF-1 on chromosome 3.**

**S4 File. Sequences co-immunoprecipitated with HIF-1 on chromosome 4.**

**S5 File. Sequences co-immunoprecipitated with HIF-1 on chromosome 5.**

**S6 File. Sequences co-immunoprecipitated with HIF-1 on chromosome X.**

**S7 File. Detailed protocol for HIF-1 chromatin immunoprecipitation (ChIP).**

**S1 Fig. HIF-1 direct targets ChIP-seq IGB signals on Chromosome 1.**

**S2 Fig. HIF-1 direct targets ChIP-seq IGB signals on Chromosome 2.**

**S3 Fig. HIF-1 direct targets ChIP-seq IGB signals on Chromosome 3.**

**S4 Fig. HIF-1 direct targets ChIP-seq IGB signals on Chromosome 4.**

**S5 Fig. HIF-1 direct targets ChIP-seq IGB signals on Chromosome 5.**

**S6 Fig. HIF-1 direct targets ChIP-seq IGB signals on Chromosome X.**

**S7 Fig. Quality scores of fastq reads for the input and ChIP DNA samples.**

